# Allelic spectrum of the RNA guided CRISPR/Cas9 DNA repair events at PAM associated trinucleotide repeat (NGG)n in *Caenorhabditis elegans.*

**DOI:** 10.1101/062489

**Authors:** Wadim Kapulkin

## Abstract

This work describes the experience with implementation of *Streptococcus pyogenes* Cas9 nuclease, expressed in *C. elegans* germline. The described work utilizes guide RNA-unc-22-1000 (GGAGAAGGAGGCGGTGCTGG) designed to target the polyglycine encoding stretch within the *unc-22*gene embedding the impure trinucleotide (NGG)n PAM repeat region. We describe the allelic spectrum of mutational events identified at position specified by gRNA-unc-22-1000. Of above experiments we conclude: i. the trinucleotide (NGG)n PAM repeat is a receptive target for the CRISPR/Cas9 experiments in *C. elegans* ii. we conclude the allelic spectrum indicates the gRNA-unc-22-1000 induces fairly frequent NHEJ joining events involving deletions and indels but also, a phenotypically distinct class of small in-frame deletions indicative for microhomology-mediated end-joining (MMEJ) as a result of *S. pyogenes* Cas9 activity at the trinucleotide (NGG)n PAM repeat region. We demonstrate guide RNA-unc-22-1000 could be used to modify complex transgenic *C. elegans* line expressing human beta-amyloid protein. We provide the evidence for bi-allelic transaction resulting from Cas9 action recovered in experiment in CB1138 *him-6* background. We contrast the expected performance of gRNA-unc-22-1000 with guide targeting another type of *C. elegans* repeat embedding *S. pyogenes* PAM, the telomeric repeat (TTAGGC)n. We propose that preferential frame restoring MMEJ repair of the Cas9 cut at the 'modules' encoding for poly-glycine in +2(NGG)n position could be useful mode of genome engineering at the naturally occurring (NGG)n PAM embedding repeats dispersed across animal genomes.

## Introduction

Clustered Regularly Interspaced Short Palindromic Repeats (CRISPR) is regarded as a class of adaptive immunity system protecting bacteria and archaea against phages and some other invading genetic elements (Deltcheva et all 2011). CRISPR system consist of ribonucleoprotein complexes and uses repeat/spacer short RNAs along with Cas (CRISPR associated proteins). The two-component system of single guide RNA (sgRNA) directing the Cas9 nuclease, operating in *Streptococcus pyogenes*, with the specificity dictated by twenty 5' base pairs of guide RNA (Jinek et al. 2012) has been proven sufficient to cleave target double-stranded DNA in *C.elegans* chromosomes (Friedland et al. 2013). In *C.elegans* Cas9-CRISPR induced lesions were reported to occur at reasonable frequency to facilitate loss-of-function studies and proposed as mediator for the transgene instructed repair procedures (Plasterk et.al. 1992, Dickinson et al. 2013). Regardless of the optimism however, still individual sgRNAs of the desired specificity, differ significantly in their biological properties, with some potent but so far poorly explained guide intrinsic and or guide extrinsic effects (i.e. observed in some synthetic RNA guides for example particularly poorly tolerated as transgenes, resulting in profound embryotoxicity and or lethality) essentially restricting their efficient *in vivo* use.

Provided the requirements for the efficient *in vivo S. pyogenes* Cas9 cleavage, here we investigated design and synthesis of in vitro modified gRNA genes at the unusual region embedded into locus of *C. elegans* encoding for UNC-22 (Moerman et al. 1979, Moerman 1980, Moerman et al. 1984, - 1986, - 1988). The unusual region consist of (NGG) impure trinucleotide repeat encoding for stretch of glycines present in UNC-22.

Given *S. pyogenes* Cas9, requires the Protospacer Adjacent Motif (NGG), for activity, we hypothesized repeats encoding for polyglycines could prove preferential targets for *in vivo* DNA cleavage. Below we demonstrate the RNA-guided Cas9 activity, resulting in removal of the polyglycine encoding trinucleotide stretch from *unc-22* coding sequence results in variably expressive distinct twitchin phenotype.

## Materials and methods

### Timeline

The experiments were conducted starting from 2013 fall and finished Jan 2015 (as the stipend awarded by Max Planck Institute CBG, has come to the expiry). The draft of manuscript were written starting from Dec 2013, and communicated to immediate coworkers via <google-docs>, followed by abbreviated communication submitted to Worm Breeders Gazette (Kapulkin et al. Aug 2014). Final versions of the manuscript were prepared in 2015-2016 in Warsaw, Poland.

### Primers

M13F and M13R primers were used to amplify U6 promoter driven klp-12 encoding sgRNA 379-1256bp segment (www.addgene.org/46170/). Internal primers sgRNA:U6--unc-22-F and sgRNA:U6--unc-22R were used to replace klp-12 targeting segment of above vector, with guide targeting eight exon of ZK617.1 (chrIV:11993024-11993044 GGAGAAGGAGGCGGTGCTGG). Primers targeting C. elegans telomeric repeat (TTAGGC)n were designed accordingly.

sgRNA:U6--unc-22 - F 5'- ggagaaggaggcggtgctgg gttttagagctagaaatagc

sgRNA:U6--unc-22 - R 5'- ccagcaccgcctccttctcc aaacatttagatttgcaattc

sgRNA:U6--TEL -F 5'- ggcttaggcttaggcttaggctt gttttagagctagaaatagc

sgRNA:U6--TEL-R 5'- aagcctaagcctaagcctaagcc aaacatttagatttgcaattc

were used for PCR synthesis of guide RNAs.

unc-22-seqF 5'- tttctctgaccacaatgcttc

unc-22-seqR 5'- tttcaatgagtaactttctgc

were ‘diagnostic’ primers amplifying the target (NGG)n repeat associated PAM with ~200bp flanks on each side.

sqt-1-F 5’- tatggagtactgtagttccta

sqt-1-R 5’- gaatgtttcaacatttgccaa

were used to amplify sqt-1 allele (sc13). The sqt-1(sc13) causes a left Roller phenotype that can suppress the rol-6(su1006) right Roller phenotype of CL4176, and was used as a transformation marker in some experiments (along with pCFJ90 or pPD122.36).

### Plasmids

#### Cas9 constructs

Peft-3::cas9-SV40_NLS::tbb-2 3'UTR containing codon optimized Cas9_SV40 NLS with intron inserted at position 3113-3163, and SV40 NLS inserted from position 5192-5227 (http://www.addgene.org/crispr/calarco/). The above plasmid use eft-3 (modified from Mos1 construct procedures developed by the Jorgensen Lab (Frøkjær-Jensen et al. 2012) promoter to drive germinal expression of the above construct. Similar eft-3 promoter construct pDD162 (Dickinson et al. 2013) including empty gRNA gene on the same plasmid were used in some experiments.

#### sgRNA constructs

PCR synthesized U6 promoter driven sgRNA-unc-22-1000 introduced (GGAGAAGGAGGCGGTGCTGG), were followed by PCR product amplified constant region of sgRNA and 3’ utr (Fig. 1A.), were inserted into vaccinia virus topoisomerase vector (Invitrogen). Background colonies from original vector (*bla*) were counterselected with kanamycin.

pJA58 *dpy-10* plasmid (addgene.com) and cn64 repair template OH-5’3’ oligo-deoxyribonucleotide (supplied by www.biomers.de) according to (Aribere et al. 2015).

**Fig. 1.**
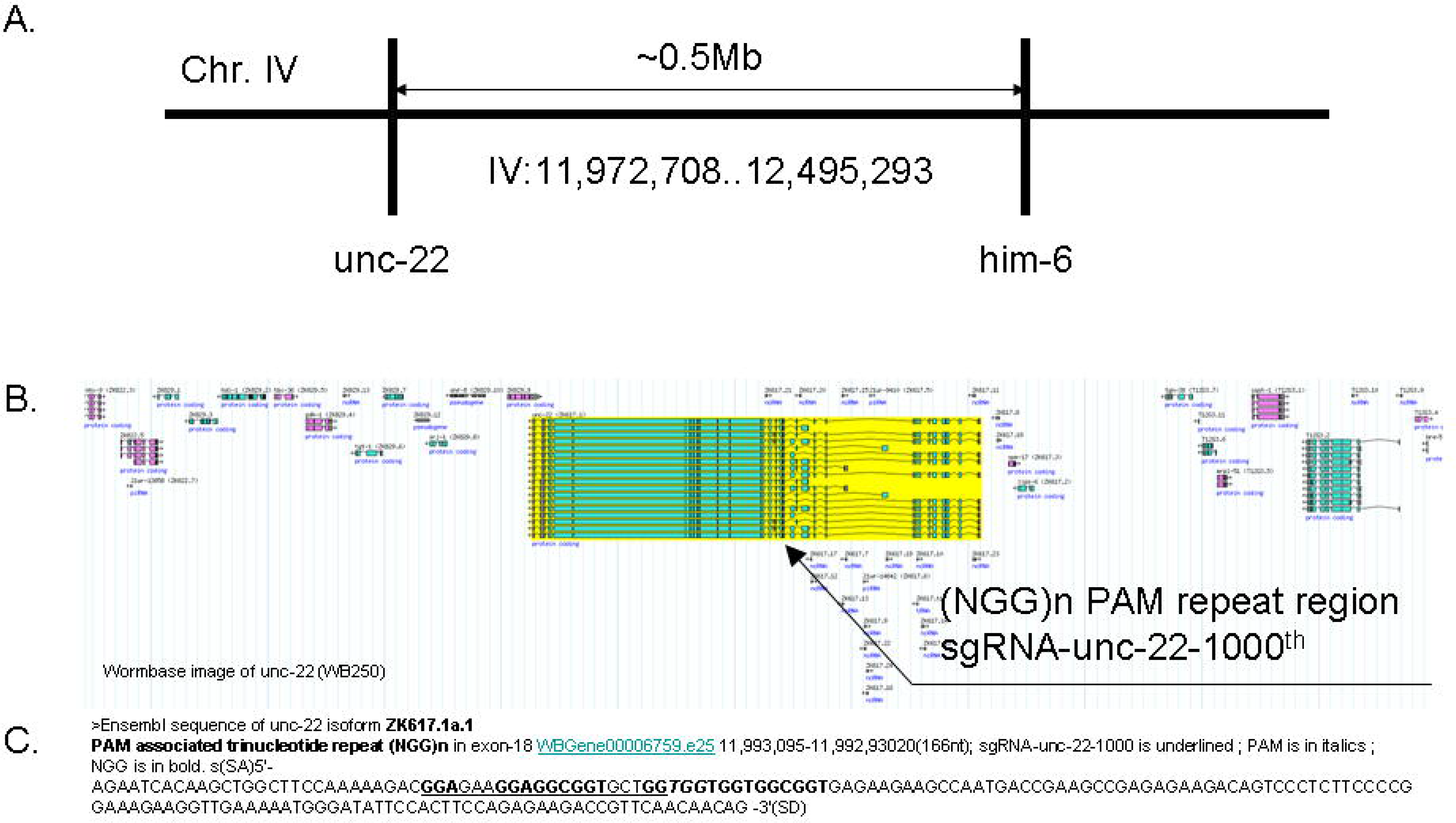
The chromosomal landscape of unc-22 IV. A. Ideogram of the interval of the chr.IV, specifying the genetic distance between unc-22 and him-6. B. Wormbase (WB250) image of the unc-22 locus. Neighboring genes are shown to demonstrate alternative transcript isoforms complexity of unc-22 (black arrow indicates approximate position of sgRNA-unc-22-1000^th^). C. Sequence of the exon 18^th^, embedding the (NGG)n repeat.

### *C. elegans* strains and transgenic modifications

The particular experiments are summarized in the Table 1. CB1138 *him-6(e1104)* (Hodgkin et al. 1979), *smg-1(cc546ts),dvIs27* CL4176 (Link et al. 2003), *unc-119 (ed3*) DP38 (Maduro et al. 1995), *sqt-1(sc13)* BE13 (Sytal et al. 2012), N2 Bristol strains were maintained as described by (Brenner 1974) and some strains were received from CGC (www.cgc.org).

**Table 1.**
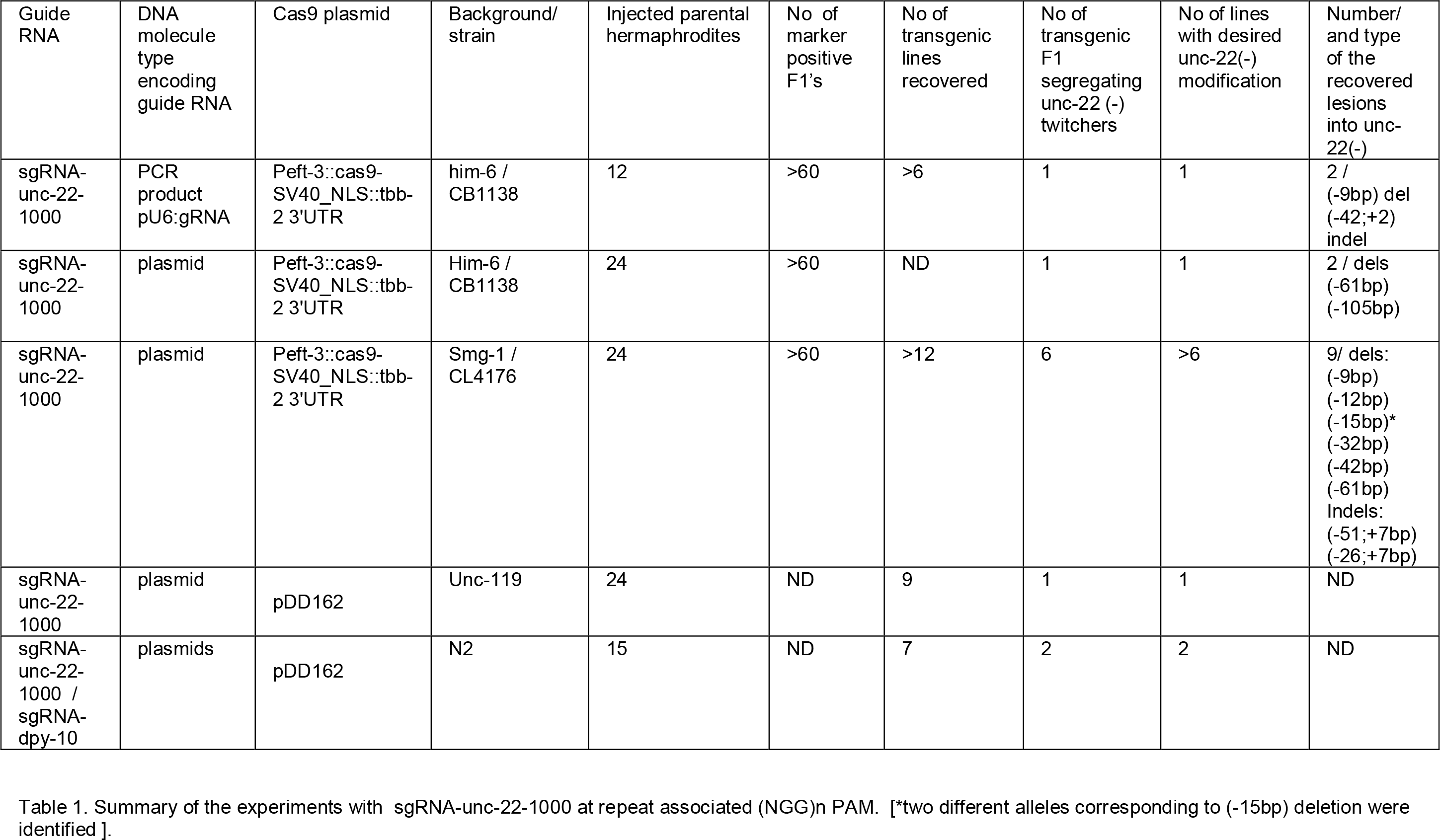
Summary of the experiments with sgRNA-unc-22-1000 at repeat associated (NGG)n PAM. [^*^two different alleles corresponding to (−15bp) deletion were identified].

| Guide RNA | DNA molecule type encoding guide RNA | Cas9 plasmid | Background/strain | Injected parental hermaphrodites | No of marker positive F1's | No of transgenic lines recovered | No of transgenic F1 segregating unc-22 (-) twitchers | No of lines with desired unc-22(-) modification | Number/ and type of the recovered lesions into unc-22(-) |
| --- | --- | --- | --- | --- | --- | --- | --- | --- | --- |
| sgRNA-unc-22-1000 | PCR product pU6:gRNA | Peft-3::cas9-SV40_NLS::tbb-2 3'UTR | him-6 / CB1138 | 12 | >60 | >6 | 1 | 1 | 2 / (-9bp) del (-42;+2) indel |
| sgRNA-unc-22-1000 | plasmid | Peft-3::cas9-SV40_NLS::tbb-2 3'UTR | Him-6 / CB1138 | 24 | >60 | ND | 1 | 1 | 2 / dels (-61bp) (-105bp) |
| sgRNA-unc-22-1000 | plasmid | Peft-3::cas9-SV40_NLS::tbb-2 3'UTR | Smg-1 / CL4176 | 24 | >60 | >12 | 6 | >6 | 9/ dels: (-9bp) (-12bp) (-15bp)* (-32bp) (-42bp) (-61bp) Indels: (-51;+7bp) (-26;+7bp) |
| sgRNA-unc-22-1000 | plasmid | pDD162 | Unc-119 | 24 | ND | 9 | 1 | 1 | ND |
| sgRNA-unc-22-1000 / sgRNA-dpy-10 | plasmids | pDD162 | N2 | 15 | ND | 7 | 2 | 2 | ND |

Transgenic modifications were carried according to the protocols established in the field (unless stated otherwise – where PCR amplified DNA was spin dialysed over the Millipore column, than filtered over 0.45um membrane). Briefly the Cas9 encoding, sgRNA encoding plasmids or other plasmids used as selectable markers (or equivalent PCRed material), were injected into hermaphrodite gonads as column purified material. Prior to injection *C. elegans* young adult hermaphrodites were kept at 16°C (well fed over at least two generations prior to injection procedure).

### Mutation screening and sequencing

Candidate unc-22(−) animals were picked out of F2 plates (except for one case were F1 were identified), allowed to self and progeny were used directly for diagnostic PCR. The remaining plates were supplemented with 0.1mM Levamisole (or equivalent strength of Tetramisole) (Lewis et al. 1980), and re-scored to ensure all CRISPR/Cas9 mutations were recovered. ‘Diagnostic’ PCR products, on candidate unc-22(−) lines were resolved on 1.5% agarose gel. Larger deletions (>25bp) were apparent directly on the gel as faster migrating bands, and all products were confirmed by standard Sanger sequencing at MPI-CBG sequencing facility. Sequences were aligned manually and with / in Genious software.

## Results

### Initial experiments with PCR modified guide RNA genes : efficient modification the PAM embedded trinucleotide repeat associated specified by sgRNA-UNC-22-1000

The single guide RNA was designed, predicted to target (NGG)n PAM repeat region within exonic segment of the C. elegans *unc-22* gene. C. elegans *unc-22* gene is fairly large gene encoded by at least 30 exons, with apparent and often complex alternative splicing patterns (quoting the wormbase information on UNC-22: ‘unc-22 encodes twitchin, a giant intracellular protein with multiple fibronectin-and immunoglobulin-like domains and a single protein kinase domain that is homologous to titin (OMIM:188840). UNC-22 is required in muscle for regulation of the actomyosin contraction-relaxation cycle and for maintenance of normal muscle morphology; UNC-22 associates with myosin and is localized to A-bands; in vitro, UNC-22 can phosphorylate myosin light-chain peptides and can undergo autophosphorylation’) and still certain of the class of <u>unc</u>oordinated alleles including unc-22 alleles are resistant to the anti-helmic drug levamisole, according to prototypical method for their selection first described by (Brener 1974). *unc-22* was chosen for our experiments because of the distinct and unique phenotype, reported in earliest classical genetics screens (Brenner 74) and since that been subject of number of reverse genetic approaches, including Tc1 transposon tagging (Moerman et al. 1986) and RNA interference (Fire et al. 1998). Of total 1609 predicted available guide RNA candidate guide sequences matching the pattern 5'G-N19-NGG-3' predicted (where …-NGG-3’ denotes PAM consensus required for *S. pyogenes* Cas9 <genome.ucsc.edu/cgi-bin/hgTrackUi?hgsid=356706829&c=chrIV&g=ct_SpCas9Cetargets_7308>) in ZK617.1 locus at chromosome IV, we selected single guide within anti-parallel strand of non-alternative exon 18 of the *unc-22* isoform 1a (Figure 1. and 4.). The unc-22 sgRNA-1000 was selected as it occurred within the impure (NGG)n trinucleotide repeat region, expected to embed the tandem repeat PAM motif otherwise known as required for the Cas9 activity. The search was focused on candidate guide RNAs with PAMs overlapping with glycine (+2) encoding repeats (NGG). As a result of an implemention of the above criteria [by combination of the output of the mfold program http://unafold.rna.albany.edu (Fig.3) and the manual inspection], designed search returned a single region of the unc-22 gene were sgRNA guide (GGAGAAGGAGGCGGTGCTGG) was chosen to target the *unc-22* locus at the trinucleotide PAM repeat region. We termed *unc-22*-sgRNA-1000, since it was the 1000th sgRNA of those predicted to lay within the *unc-22* gene. (Suppl. Table 1.).

We initially choose to manipulate the sgRNA genes encoded by pU6:unc-22-sgRNA-1000, via PCR. This was done due to the following rationale: PCR fused segments of DNA (Hobert 2002) could be obtained from routine in vitro reactions, and obtained fused-PCR products could be tested directly for their biological activity in vivo by injecting into hermaphrodite gonads (prior to attempting with construction of plasmids in *E. coli*). The above PCR-based strategy was designed and implemented to modify the specificity dictating region of guide RNA genes as we suspected it might be necessary test several candidate guide RNAs prior to identifying the one with sufficient biological activity to trigger the observable *in vivo* effects of Cas9 action within the specified region of unc-22 gene (located in *C. elegans* chromosome IV, Fig. 1). In the initial attempt to test for the efficient CAS-9 activity, in *C. elegans* germline, PCR synthesized material encoding for the modified sgRNA genes was co-injected along with established plasmids (encoding for eft-3 promoter driven Cas9 see materials and methods) protein and reporter markers, into 12 hermaphrodites of *him-6(e1104)* genotype (Hodgkin et al. 1979). In this initial experiments we intended to compare the biological effects of two PCR products representing the modified at the 5’ specificity-dictating regions of two different guide RNA genes. The first PCR product was the pU6:unc-22-sgRNA-1000. The second PCR product was the pU6:TEL-sgRNA specifying *C. elegans* telomeric repeat (TTAGGC)n. (The second PCR product was intended to induce CAS-9 cleavages in multiple regions corresponding to *C. elegans* telomeric repeats present in all six chromosomes and was regarded as negative control in the experiment). [See Supplement materials: *C. elegans* telomeric repeats are tandem hexa-nucleotide repeats embedding NGG PAM pattern, dispersed at the ends of every six of the *C. elegans* chromosomes. Given the designed guide RNA specifying telomeric repeat embedding *S. pyogenes* Cas9 5'-AGG-3' pattern in a form of non-tandem PAM repeats is present in the *C.elegans* genome sequence at least 700 times (www.wormbase.org) this multivalent guide RNA is expected to induce CAS-9 cleavage at the telomeric and sub-telomeric regions of every chromosome). We expected that by provoking presumably massive on-target CAS-9 activity at the multiple chromosomal locations, specified by telomere embedded 5’-AGG-3’ repeat PAM, will effect in the intense cellular repair response and induce numerous blunt-ended double stranded brakes into every of the nuclear DNA molecules. Provided the desired, heritable DNA modification resulting from the CAS-9 generated DNA cleavages, have to involve the chromosomes present in the germ-line cells, we expected the multivalent guide RNA specifying the telomeric repeat 5’-AGG-3’ repeat PAM region would cause the excessive DNA damage and induce other negative (toxic) effects directly into gametes.] Indeed, when we initially compared the overall efficiency of the transformation of the hermaphrodites injected with two above sgRNA guides specifying the UNC-22 sgRNA-1000 (NGG)n glycine encoding trinucleotide PAM associated repeat and sgRNA-TEL (TTA GGC)n telomeric PAM repeat, we have observed the lethality or sterility in transformed F1s resulting in drastic (estimated at order to two orders of magnitude) drop in the number of stable transgenic lines we recovered. In the first 12 injected hermaphrodites, with above modified UNC-22 sgRNA-1000 (NGG)n trinucleotide PAM repeat, transgenic lines (as judged by transmission of co-injected transformation markers) emerged rapidly and the desired genotype of unc-22 animals was identified within the first six extrachromosomal array transmitting transgenic lines. In contrast injection of the second PCR-fused product (different from sgRNA-unc-22-1000 in only 20 nt specifying the 5’ end of guide RNA substituted to telomeric repeat 5’-AGG-3’ non-tandem repeat PAM) diluted over the same preparation of the co-injection transgenic marker sample mix, resulted in only single array transmitting line recovered of an entire transgenic descent of all 12 injected animals. From this initial experiment we have concluded the PCR products could be used readily to alter the specificity of the 5’ends of synthetic - guide RNA genes and that the control construct (TTA GGC)n telomeric PAM repeat specifying the 5’ end of guide RNA pose the significant threat to procedures requiring the transgenic lines by inducing lethal effects. While cautious not to over-interpret the above negative results, having in mind the presently uncertain nature of the observed unwanted lethal or embryo-toxic effects we emphasize the developed pilot procedure of injecting the PCR-stiched guide RNA genes helps to identify sub-lethal or otherwise inefficient synthetic 5’CAAA3’ RNA guides, early in the experiments prior to attempting to establish first clones of bacterial replicons propagating modified guide RNA genes. We envision that complex libraries of the parallel modified PCR products encoding guide RNAs specifying for any given genomic region could be conveniently PCR synthesized and tested directly by injecting into hermaphrodite gonads, to screen for the most optimal variants present in the initial mixtures of the predicted pool of candidates (see discussion).

The initial experiments described above conducted with PCR modified sgRNA genes, were conducted using *him-6(e1104)* CB1138 background. *C. elegans him-6* is the orthologue of a human (and mouse (Luo et al. 2000)) BLM encoding gene. BLM is a RecQ DNA helicase and is mutated in the Bloom syndrome (0MIM:604610). Bloom syndrome is a form of progeria with unusual segregation of sister chromatids during the cell division. In *C. elegans* Blm orthologue *him-6* (Hodgkin et al. 1979) is located on IV chromosome and *him-6* is linked with *unc-22* gene on the *C. elegans* chromosome IV (ZK617.1 / unc-22 is separated from T04A11.6 / him-6 on IV:5.45802 to IV:5.98937 converting to roughly 0.5314cM (wormbase220) Fig.1.). We took an advantage of this genomic arrangement and have established in the CRISPR/Cas9 experiment, that lesions into the UNC-22 trinucleotide PAM repeat region indeed result in the identification of the doubly homozygous *him-6(−);unc-22(−) IV* genotype (linked on chromosome IV and separated by approximately only half of the megabase). Encouraged by the initial positive results with PCR-fused guide RNA gene we prepared a plasmid clone containing the fused-PCR modified sgRNA-UNC-22-1000 gene driven by U6 promoter. In order to test weather the initial results of injecting the low concentrated PCR modified guide RNA gene would result in consistent alternation to the *unc-22* trinuclotide PAM repeat glycine encoding region, the purified material derived from *E. coli* plasmid replicons (containing PCR modified guide RNA gene, sgRNA-UNC-22-1000), was injected to gonads of 24 *him-6*(CB1138) hermaphrodites. In this experiment *him-6* animals were injected with sgRNA-UNC-22-1000 plasmid, diluted over the same injection mix as in initial experiment (however plasmid DNA was included at 5-10 fold increased concentration, when compared to the estimated concentrations of the raw PCR product). In this particular experiment one of the injected *him-6* hermaphrodites, segregated the exceptional doubly homozygous animal of <u>*him-6; unc-22(−)* genotype in the F1</u>. Provided the characteristic twitching phenotype prominent and apparent in the F1 recovered from the experiment (consistent with homozygous disruption of the *unc-22 IV*), inherits as a recessive trait, we have reasoned that this exceptional unc-22(−) F1 hermaphrodite, most likely had both sperm – and egg-derived paternal chromosomes modified as a consequence of the Cas9 cleavage. We note however, that an alternative explanation does exist, given the unusual segregation of non-disjunctional gametes were reported in *him-6* (Hodgkin et al. 1971). In particular (Haack & Hodgkin 1991) established that observable fraction of progeny represents the genotype concordant with uniparental disomy at every *C. elegans* linkage group (see discussion) and thereof both Cas9 modified chromosomes IV we recovered could represent the maternal line of descent.

We have established, the injection of the sgRNA-UNC-22-1000 with Cas9 encoding plasmid, into CB1138 *him-6* line, yielded consistently the progeny animals of the expected phenotype, we included into ours experiments the control group of *him-6(+)*. For the clarity the control *him-6(+)* animals were injected with the same preparation of mixture of CRISPR/Cas9 plasmid DNA, as in the above experiment on *him-6(−)* genotype. For the purpose of this control experiment we have selected the *him-6(+)* line CL4176 (Link et al. 2003). This was dome for the following reasons: i. CL4176 carries homozygous, X integrated transgenic array, and is maintained as isogenic populations consisting of hermaphrodites, with males occurring at frequencies comparable to wild type prototype N2 (~0.02%). ii *dvIs27* is X integrated transgenic arrays, contributing the expression of the transgenic variant of collagen *rol-6(d)* (su1006) and beta-amyloid encoding transgene (pAF27) into *C. elegans* muscle cells. iii. CL4176 line carries also a temperature-sensitive mutation in the SMG-1 encoding gene. A temperature-sensitive mutation *smg-1(cc546ts)* was developed by the Fire lab as a means to engineer conditional transgene expression (WBG Getz et al. 1997). *smg-1(cc546ts)* background might contribute to the overall beta-amyloid phenotype observed in CL4176 (*dvIs27*) and disruption to *unc-22* functions in the above background greatly aggravates the uncoordinated phenotype observed in ‘twitchers’ (Kapulkin et al. 2005, and Kapulkin WJ unpublished observations). We note that in the background we have chosen, neither *dvIs27* on chr. X nor *smg-1(ts)* on chr. I are linked to *unc-22* on chr, IV and therefore are easily removed by crossing with WT males.

Collectively we reasoned that by attempting to modify with the CRISPR/CAS-9 system the complex transgenic line (where transgene expression from X chromosome locus is controlled by the unlinked homozygous temeperature-sensitive allele into locus of SMG-1 on chromosome I), we could possibly recover new *unc-22* variants at the improved rate. This in turn would be relevant route if candidate (i.e. bata-amyloid toxicity suppressors (Lopez 2009)) mutations would be to be introduced with CRISPR/Cas9 into CL4176. Upon injection of the CRISPR/Cas9 components into gonads of 24 hermaphrodites (again using exactly the same preparation of the injection mixture as injected into *him-6(−)* hermaphrodites), we immediately identified at least three marker positive F1 animals (of sixty initially singled transgenic F1) of two independently injected parental animals segregated severely uncoordinated F2 twitchers. Observed sgRNA-unc-22-1000 guided CRISPR/CAS-9 induced unc-22 phenotype in CL4176 line, manifested in populations of animals where frequent twitches are imposed onto almost entirely non-mobile bodies. We termed those entirely uncoordinated *unc-22(−)* ‘spastic twitchers’. Spastic twitchers in CL4176 background are strikingly apparent and different in overall phenotypic severity from *unc-22* alleles maintained in the wild type background (i.e. *e66* reference twitchers described by classically by Brenner). Disruptions into *unc-22* in wild type or *him-6 (−)* background are described as myoclonic spasms imposed onto otherwise normal movement. Spastic twitchers induced by CRISPR/Cas9 into CL4176 in contrast appear as strikingly apparent (in the absence of the levamisole selection) and almost entirely immobile (Supplement Movie 1).

While screening for the CRISPR/Cas9 modified animals we also noted that at least three populations segregated considerably weaker *unc-22(−)* animals (weaker when compared to *e66* reference twitchers described in classical Brenner’s screens). Those weak alleles were identified, amongst F2-F3 populations born from of at least three other array transmitting transgenic F1’s (transgenic siblings of ‘spastic twitchers’ were borne from same injected parental animal). Above ‘weak’ *unc-22(−)* siblings were noted later in the experiment, based on the occasional muscle spasms, infrequently affecting the normal movement.

Amongst *unc-22(−)* disrupted, transgenic array transmitting CL4176 hermaphrodites, we recovered two sibling twitcher lines, growing considerably slower than the other lines. Remarkably the two of the *unc-22(−)* transgenic arraray transmitting lines apparently represented homozygous insertions of the extrachromosomal arrays (containing the mixture of injected plasmids encoding for sgRNA-unc-22-1000, Cas9 and co-segregating plasmids encoding for transformation markers). The two above integrated lines were mated with males to establish the possible linkage between integrated arrays and *unc-22(−)* on IV, (and to eliminate the *dvIs27* background transgene integrated on X). The progeny of the cross of the first of the above integrated sgRNA-unc-22-1000 Cas9 array lines segregated expected non-rol progeny indicating for the loss of *dvIs27(−)* integrated on X. When allowed to self, integrated sgRNA-unc-22-1000 Cas9 array transmitting heterozygous non-twitching hermaphrodites segregated expectedly about ¼ of unc-22(−) progeny and co-segregated integrated Cas9 encoding transgenic array. (the other integrated Cas9 transgenic array line failed to segregate expected non-rol F2.) This might suggest above integrated Cas9 transgenic array lines might represent integration events who occurred independently and perhaps concomitantly to gRNA-unc-22-1000 specified effects of Cas9. This observation might be relevant as suggests that background integration events of Cas9 encoding arrays might persist in experiments with CRISPR/Cas9 system in *C. elegans*.

### The recurrent pattern of *S. pyogenes* Cas9 induced DNA lesions in regions specified by the repeat associated (NGG)n PAM: the evidence suggesting involvement of an alternative repeat repair pathway MMEJ

Provided the repetitive nature of the UNC-22 region specified by the guide RNA-unc-22-1000, we asked the question regarding the molecular pattern of the DNA lesions, presumably present in the him-6(+) and him-6(−), unc-22(−) twitcher lines. *unc-22* is a large gene, predicted to encode for muscle protein with multiple immunoglobulin-like domains as well as kinase domain at C-terminus. The (NGG)n repeat region we choose to specify 3’ PAM repeat in the guide RNA sequence, is embedded between third and fourth Ig-like domain. The (NGG)n trinucleotide repeat region is encoded by the small exon and translates into impure poly-glycine-repeat peptide, where stretch of glycines is interrupted by single alanine residue (<u>GlyGlyGly</u>Ala<u>GlyGlyGlyGlyGly</u>). To test for the Cas9 effects specified by the guide RNA-unc-22-1000 (predicted to specify PAM in the above +2(NGG)n poly-glycine encoding repeat), the isolated unc-22(−) clonal lines were allowed to self, and products of PCR amplification on DNAs were sequenced along with control, non-twitching sibling lines derived from same, injected hermaphrodite. The sequencing results demonstrated lesions ranging from 9bp to 105 base pairs removed, altering or entirely removing poly-glycine encoding repeat (Fig.2). The details specifying deletion breakpoints are listed with (Table 2.) Based on the phenotypic assessment of the severity of unc-22 twitcher lines, we observed the following clustering of the recovered (respectively stronger and weaker than *e66*) unc-22 alleles. Strong unc-22 alleles tend to remove entire poly-glycine encoding repeat with adjacent flanking exon sequences by either longer frame restoring deletion (−42bp), frame altering deletions (−61bp) and (− 32bp) or frame altering indels (−51;+7bp), (−42;+2bp) and (−26;+7bp). The largest (− 105bp) deletion, recovered from the exceptional *him-6(−); unc-22(−)* F1 removes the entire 5’ part of the exon encoding for PAM associated trinucleotide repeat, including 3’ splice site and presumably alters the splicing pattern unc-22 (Fig.2, data not shown). In contrast, unc-22 alleles classified as ‘weaker than *e66’* represent entirely, small frame restoring deletions (−9bp), (−12bp) and (−15bp) (Table 2.). Weak alleles were found to remove the codons for alanine and two, three or four adjacent glycines (respectively), and convert the impure (NGG)n PAM associated repeat into pure poly-glycine encoding repeat (converting <u>GlyGlyGly</u>Ala<u>GlyGlyGlyGlyGly</u> into hexa-glycine, penta-glycine or tetra-glycine encoding repeat (Fig. 4C.)). In a matter of the fact phenotypic manifestations observed in homozygotes for weaker alleles identified in the study, could be described as subtle. Animals locomotion were affected by twitches and spasms in muscle cells in only occasional manner and otherwise barely noted. In ours hands it is the sensitized background of integrated *dvIs27* array line *smg-1(ts)*, where improvement in recovery of the small frame restoring changes is observed. In this sensitized background, approximately half of the recovered alleles at (NGG)n trinucleotide PAM embedding repeat region were identified as small (−9bp) (−12bp) (−15bp) frame restoring deletions (Fig.2. Aligment of the alleles identified at PAM repeat region and Table 1. and Table 2.). Otherwise, when recovering from him-6(−) experiments with both PCR-fused and plasmid encoded sgRNA-unc-22-1000 gene only single event of 9bp deletions were recorded. Provided the repetitive nature of the (NGG)n PAM containing region specified by guide RNA the (−9bp) deletion could be aligned in five possible positions. Similarly (− 12bp) deletion could be aligned in three possible position. We note this repetitive attribute of (NGG)n might complicate the possible interpretation concerning the repair events involved in solving of an initial Cas9 cut. Given the several observed independent (9bp) and (12bp) deletions we could not exclude if those arose from cut/repair events at the specified position or if those could result from use of an alternative adjacent NGG PAM. Provided the repetitive nature of the trinucleotide (NGG)n associated PAM, we propose the alternative, the repeat repair pathway referred as MMEJ (microhomology-mediated end joining) is used instead of NHEJ (non-homologous end joining) to repair the sgRNA-unc-22-1000 specified Cas9 cut (see discussion).

**Fig. 2.**
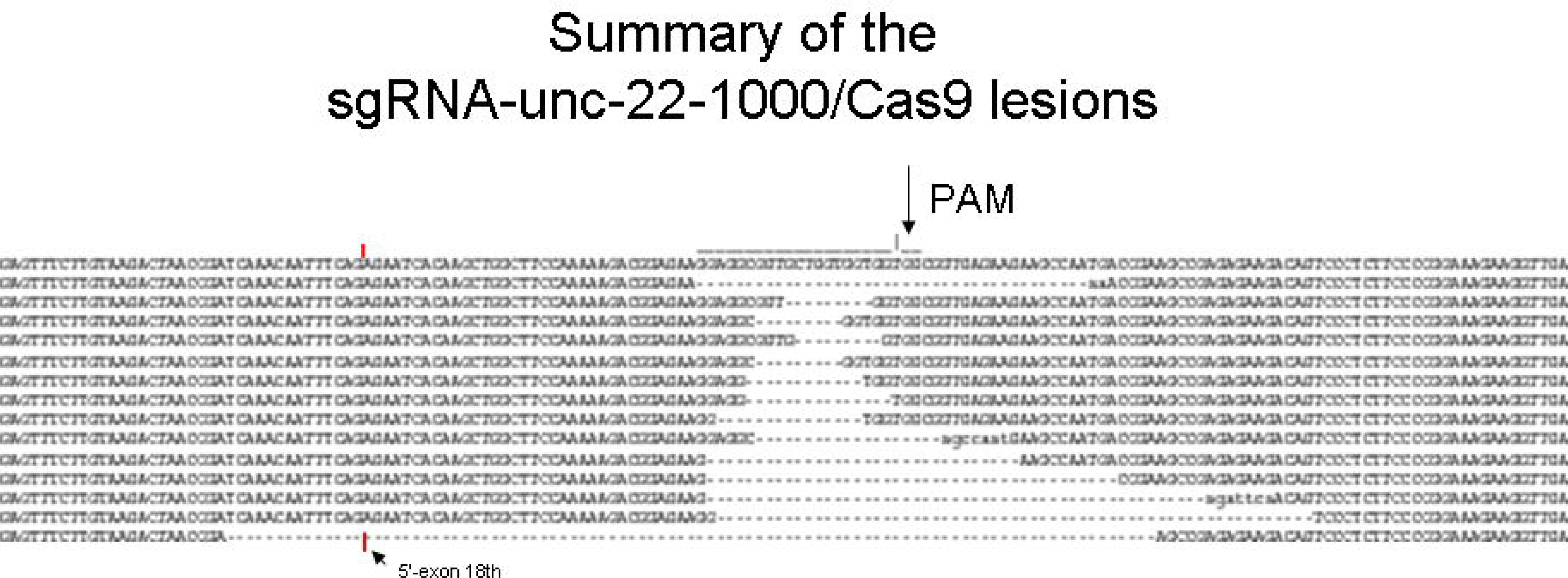
RNA guided lesions induced by Cas9 at exon 18 of unc-22 aligned to WT (top line, overlined is sgRNA-unc-22-1000 sequence, followed by (TGG) PAM), in the following order sg1000-(−42+2); sg1000-(−9)a; sg1000-(−9)b; sg1000-(−9)c; sg1000-(−9)d; sg1000-(−12); sg1000-pU6-(−15)a; sg1000-(−15)b; sg1000-(−26+7); sg1000-(−32); sg1000-(−42); sg1000-(−51+7); sg1000-(−61), sg1000-(−105). Legend:(−) demarks missing base(n) lowercase marks inserted base. Notes: 1. sg1000-(−9)a-d were isolated from independently injected parental hermaphrodites (sg1000-(−9)a-d are aligned in four different positions over the WT) 2. sg1000-(−105) is predicted to remove 5’exon-intron junction (is marked by the red line/black arrowhead. Exon 18^th^ starts as 5’-AGAATCACAAGCTGG-…−3’ in the above picture)

**Fig. 3.**
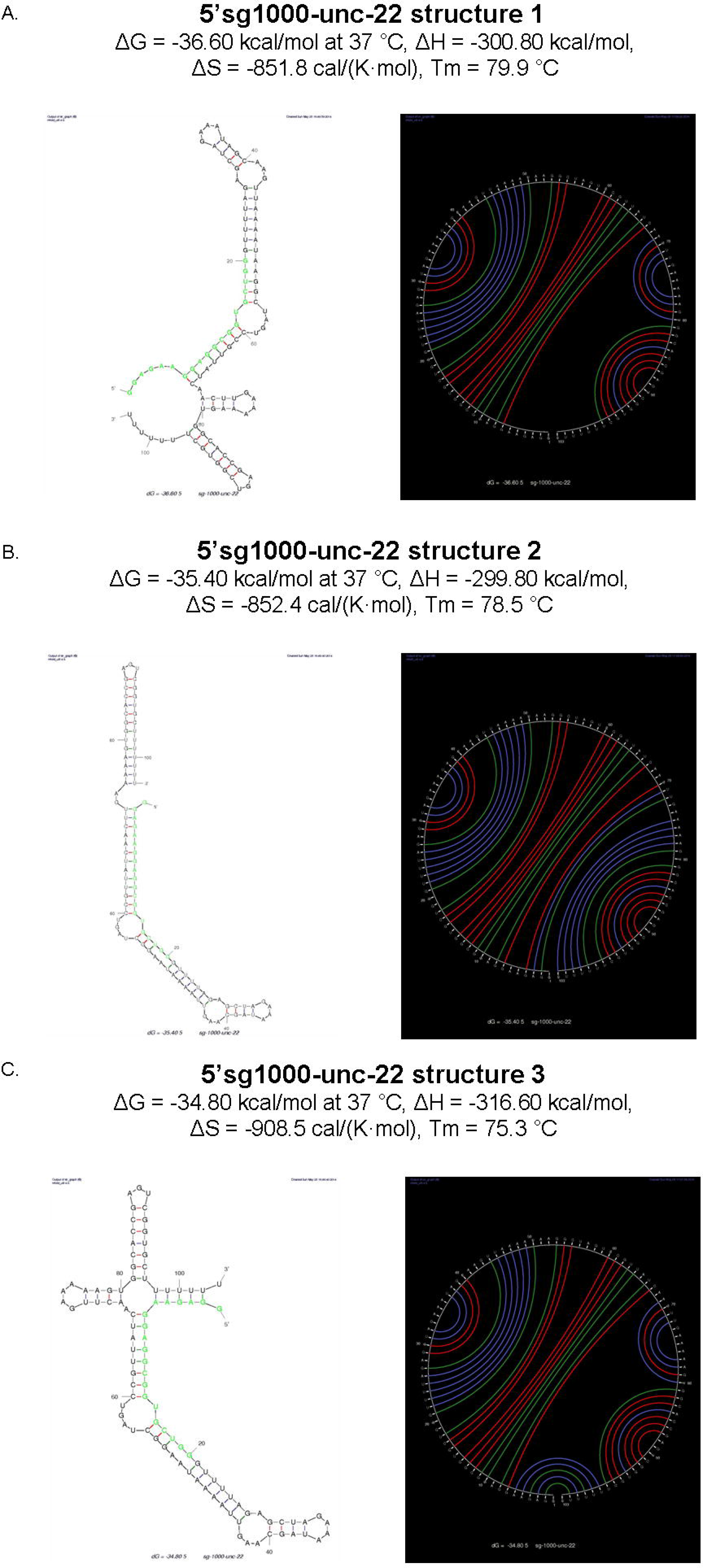
Three individual foldings of the sgRNA-unc-22-1000 predicted by the mfold program (A., B., C,), [<http://www.albany.edu/rna/withMF0LD2.3> with following parameters; Tm (°C) assuming a 2 state model at 37 °C, linear RNA folding,Ionic conditions; [Na+] = 1,0 M, [Mg++] = 0,0 M, (where standard errors are roughly ±5%, ±10%, ±11% and 2-4 °C for free energy, enthalpy, entropy and Tm, respectively). Note the above model assumes ‘single RNA molecule 1:1 stechiometry’,] sgRNA-unc-22-1000 th is in position 1-20 (GGAGAAGGAGGCGGUGCUGG) and bases are given in green crRNA-uric-22-1000th is in position 20-32 (GUUUUAGAGCUA), GAAA is a linker 5'-3' loop between crRNA and trans-activatingRNA is in position 31-33 bp, tracrRNA-unc-22-1000th is located in positions 33-103 at −3' end (UAGCAAGUUAAAAUAAGGCU AGUCC-GUUAUCAACUUGAAAAAGUGGCACCGAGUCGGUCUUUUUUU).

**Fig. 4.**
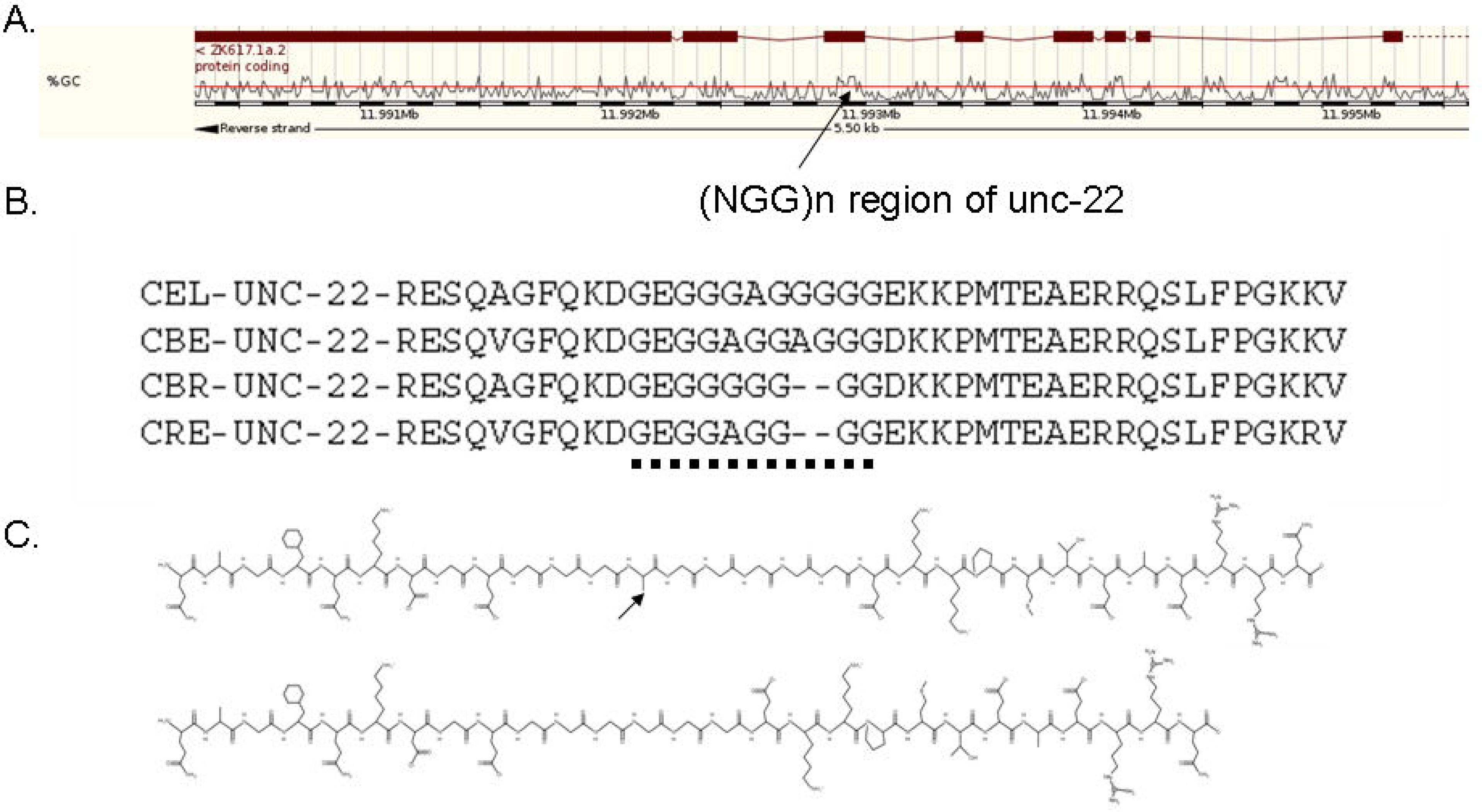
Comparative view of the (NGG)n region of unc-22. A. Ensembl image of 5.5kb around the non-alternative exon 18 (+2 polyglycine encoding stretch is indicated by arrow. Note: expected GC% peak consistent with sgRNA-unc-22-1000 predicted as 70% GC). B. Alignment of conserved (+2) polyglycine (underlined by the rectangle dot line) stretch in UNC-22. CEL-, CBE-, CBR-and CRE-denotes C. *elegans*, *C. briggsae*, *C. brenneri* and *C. ramonei* UNC-22 predicted sequence respectively. C. PepDraw images of polyglycine strech in UNC-22. Upper is WT [QAG FQ KDGE<u>GGGAGGGGG</u>EKKPMTEAERRQ], bottom is frame restoring 9bp deletion [QAGFQKDGE<u>GGGGGG</u>EKKPMTEAERRQ] resulting in weak twitcher unc-22(−) phenotype (arrow indicates the deleted alanine residue)

**Table. 2.**
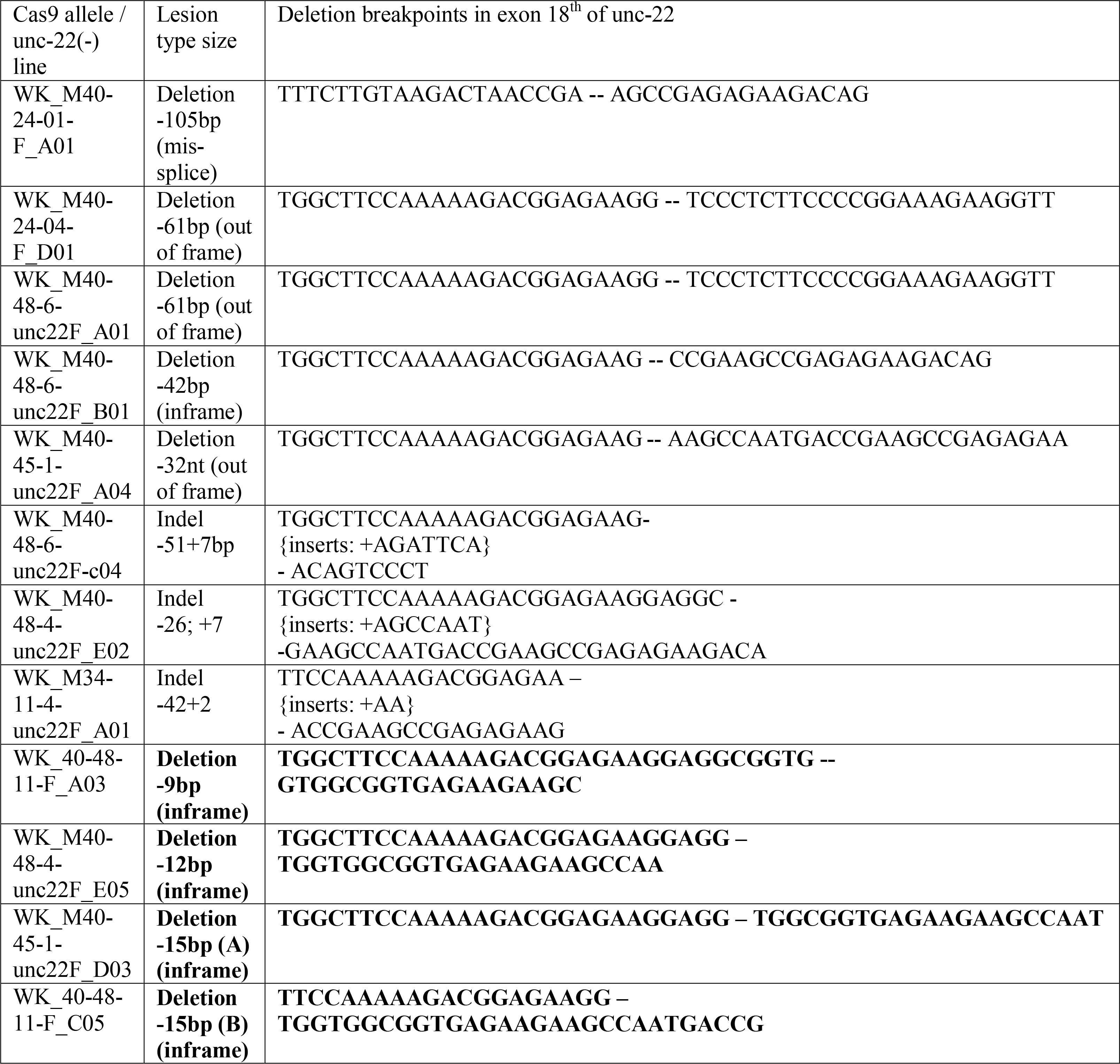
Deletion breakpoints of the sgRNA-unc-22-1000 guided Cas9 alleles in unc-22. Frame restoring deletions (−9bp) (−12bp) (−15bp) indicative for MMEJ are indicated in vbold.

Another surprising observation concerning the reoccurring SpCas9 induced deletions in independently injected animals with sgRNA-unc-22-1000, was not restricted to weak alleles representing small in-frame deletions within polyglycine repeat region (presumably repaired by MMEJ). Strikingly (−61bp) deletion, product of NHEJ repair (of strength comparable to reference *e66*), were confirmed at least twice in the homozygous F2 populations singled from progeny of at least two different independently injected parental animals. Identical (-61bp) deletions (removing the entire poly-glycine encoding repeat with adjacent 5’ and 3’ flanking sequences (Fig.2.)) were identified in the experiments conducted on him(−) and him(+) backgrounds. Of the above observation we concluded, that in the independent CRISPR/Sp-Cas9 experiments, the most prevalent lesions [in the given case of unc-22-sgRNA-1000 dictated cleavage the distinct (−61bp) deletion] apparently reoccur. We note it will be interesting to consider the possibilities altering the pattern of the *in vivo* Sp-Cas9 cleavage i.e. this might imply that the initial Cas9 cleavage, specified by unc-22-sgRNA-1000 operates under ‘some’ genomic constrain (i.e. due to the local chromatin structure or some other properties of the specified (NGG)n repeat PAM region of the unc-22 locus) and / or the DNA break repair mechanism might be perhaps biased (see discussion).

### Repeat associated (NGG)n PAM is a subject of sgRNA-unc-22-1000/CAS9 cleavage / repair in the DP38 *unc-119 (ed3)*

Having established the sgRNA-unc-22-1000, triggers the Sp-Cas9 cleavage at repeat associated (NGG)n PAM in both two tested backgrounds containing (CB1138 *him-6* and sensitized CL4176 *dvIs27*), we set to establish if we could select for the unc-22(−) mutants in *unc-119 (ed3*) (Maduro et al. 1995). DP38 line carrying *unc-119 (ed3)* allele is regarded as preferred background, in number of transgenic procedures established in *C. elegans*. DP38 background have been utilized as a transformation marker (where rescued by transgenic WT *unc-119* favors the selection of transformed animals). For example (Sarov et al. 2006) have developed the set of GFP modified fosmid-based constructs, suitable for the selection in DP38 background. We thought those fosmid-based constructs might prove the promising reagents for CRISPR-Cas9 experiments (i.e. where transformed fosmid constructs could be used as ‘transgenic rescue’ cassettes in DP38). Importantly (Dickinson et al. 2013) set up the improved CRISPR based system where *unc-119 (ed3)* background were used to select for template directed repair transactions following the Cas9 cut.

Having in mind the above considerations concerning the use of DP38, we implemented the experimental design favoring the selection of sgRNA-unc-22-1000 directed Cas9 lesions in transgenic arrays forming events by introducing the *sqt-1 (sc13)* as an auxiliary selection marker (Saytal et al. 1998). *sqt-1 (sc13)* appears as useful as transgenic marker in number of *C. elegans* backgrounds: when co-injected into to *him-6* (or other non-Rol lines) causes transformed animals to develop the left Roller phenotype, however it suppresses the dominant left Roller phenotype of *rol-6(d)*(su1006) included into *dvIs27*. We objected to test if, sgRNA-unc-22-1000 mediated effects of Cas9 at the PAM associated (NGG)n trinucleotide repeat region of *unc-22* could be selected in the *unc-119(ed3)* background using modified unc-119(+) fosmid constructs. To directly test for feasibility of this selection system in CRISPR/Cas9 experiments, we have injected mixtures of plasmids encoding for pU6:sgRNA-unc-22-1000, pelt-3:Cas9 (Dickinson et al. 2013), *sqt-1(sc13)* along with modified fosmid clones (carrying the wild type *C.br-unc-119* (Sarov et al. 2006) into gonads of 24 *unc-119 (ed3)* hermaphrodites (DP38). Expectedly we have recovered unc-119 rescued F1 animals as non-unc, *sqt-1(sc13)* rollers, who grew considerably faster than original line DP38 (this have allowed for the selection of transgenic arrays directly on the plates with injected parental *unc-119(−)* DP38 animals). We have observed that one of the first seven marker positive lines segregating the rescued progeny (homozygous for *unc-119* (DP38) recovered as Roller array transmitting lines) segregated also the expected array positive *unc-22(−)/unc-119(+)* twitchers and slow growing, array negative *unc-22(−)/unc-119(−)* double homozygotes. Provided the presence of the expected phenotype, we have initially concluded of the above experiment, that DP38 could be used as background for selecting for CRISPR-Cas9 lesions at PAM associated (NGG)n region of gene encoding for UNC-22, using the modified fosmid constructs. We note however, that when attempted to verify the integrity (NGG)n PAM repeated sgRNA-unc-22-1000 region, with standard PCR assay used in the previous experiments, we could not observe the expected amplicons (data not shown). This might be due the size of the deletion (i.e. eliminating the primer binding site), major chromosomal rearrangement (i.e. chromosome break after the double stranded Cas9 cut) or insertion of the larger DNA fragment into the initial Cas9 generated lesion or other form of the aberrant repair. Similar effects however, where Cas9 induced alleles were not accessible for PCR detection have been noted by others experimenting with CRISPR-Cas9 improvement in *C. elegans* (i.e. Arribere et al. 2014). It remains unclear if MMEJ or NHEJ events observed at (NGG)n repeat region of *unc-22* could possibly affect the repair process at PAM associated repeats. It is also presently unclear if fosmid-based repair templates, supplying the excess of the PAM embedding (NGG)n repeats incorporated into extrachromosomal arrays would remain permissive for Cas9 dependent HR transactions.

### Repeat associated (NGG)n PAM is permissive for sgRNA-unc-22-1000 specified CAS9 cleavage / repair in the presence of concomitant co-conversion with co-administrated oligonucleotide repair templates at the unlinked second site CRISPR/Cas9 target

We next set out the experiment to test if the effects of sgRNA-unc-22-1000 directed Cas9 cleavage repair could be observed at unc-22 unlinked locus. In this experiment we aimed to exclude for some unforeseen and unwanted effects associated with sgRNA-unc-22-1000 directed Cas9 cleavage at the glycine encoding PAM repeat (NGG)n region on the intended repair with co-administrated oligonucleotide repair templates. This particular experimental design was due to the failure (data not shown) of the initial attempts to observe point mutations introduced with ssDNA oligonucleotide repair reported by (Zhao et al. 2013) or fosmid-derived dsDNA repair templates, at the PAM embedding (NGG)n trinucleotide repeat region of unc-22.

We choose the well characterized oligonucleotide (Aribere et al. 2015) at *dpy-10* locus, which is efficiently repaired with oligodeoxynucleotide template converting WT N2 into distinctive *cn64* allele. Provided the *cn64* co-conversion effects discriminate between oligonucleotide repair converted F1s and background NHEJ-like events, we injected the sgRNA-unc-22-1000/Cas9 components, with plasmids providing gRNA-dpy-10 (pJA58) and 20ng/ul cn64 converting ssDNA synthetic oligonucleotide into gonads of 24 N2 hermaphrodites. Expectedly, in the F1 generation (populations where progenies of at least 15 hermaphrodites were scored positively for transgenic markers) 7 of picked rol-non-dpy animals were identified as founder populations truly converting WT *dpy-10* into *cn64*. In the second generation selected F2 progeny were screened for the recessive trait conferring to myoclonus indicative for disruption at the unc-22 twitchin gene (expectedly consistent with sgRNA-unc-22-1000 specified CAS9 mediated cleavage at (NGG)n repeat region). Indeed two of above of seven lines (Table1.) efficiently converting WT *dpy-10* into *cn64* rollers, segregated mendelian proportion of the expected F2s homozygous for *unc-22(−)*. From the above experiments we conclude: i. the concomitant *S. pyogenes* Cas9 effects on (NGG)n repeat-associated guide RNA sgRNA-unc-22-1000 locus could be selected at distant locus *dpy-10* with oligonucleotide conversion to *cn64*, ii. in the absence of the repair template supplying the glycine encoding (NGG)n repeat-associated PAM, the Cas9 effects specified by guide RNA sgRNA-unc-22-1000, do not preclude efficient templated repair on its own (which appear consistent with the perspective where the impaired templated repair at (NGG)n repeat region, does not result from generalized inability to solve the Cas9 cut / repair i.e. due to some unforeseen effects of the unusual repeat associated PAMs on the Cas9 action and or DBS repair). It is therefore expected the fraction of pre-selected *dpy-10* converted, animals will develop into double *unc-22(−)* homozygotes (leaving 5 of 7 seven in ours case, lines lacking the other second-site desired modification, specified at the unlinked unc-22 locus). This might be due to the different requirements of different guides optima or some other effects (see discussion).

## Discussion

Improved Cas9-CRISPR system developed (Jinek et al. 2013), critically relies of the ‘artifactual’ covalent 5’3’ phosphodiester bond introduced into guide RNA molecule, joining two separate CRISPR-associated small RNAs via four nucleotide (CAAA) junction loop. The 5’ prime is a ‘variable’ region dictating the specificity of derived from CRISPR RNA (also referred as crRNA) and 3’ PAM containing ‘constant’ region derived from trans-activating crRNA, to form so called single guide RNA (sgRNA). sgRNAs are of exceptional value in genetics as those if conveniently modified at 5’ end, were shown to efficiently specify Cas9 nuclease activity *in vivo* in many model experimental systems, including *C. elegans*. The designer choice of segments suitable for the modification at the 5’ end of sgRNA however is restricted to DNA segments matching the discrete consensus, referred as Protospacer Adjacent Motif (PAM). PAM sequences appear species specific and *S. pyogenes* PAM minimal consensus is a single (NGG) motif. Provided the (NGG) PAM appears strictly required for the Sp-Cas9 *in vivo* activity, we have investigated if *C. elegans* repeat-associated PAM regions (NGG)n would be suitable as the targets for sgRNA modifications.

We identified the (NGG)n repeat region embedded into coding sequence of *unc-22* and demonstrated that (NGG)n sgRNA-unc-22-1000 is sufficient to induce the DNA lesions, removing or altering the PAM. Based on the preliminary experiments with few different guide RNAs, we observed (data not shown) however that only fraction of the tested sequences performed expectedly and optimally. We therefore inspected the UNC-22 (NGG)n repeat regions for candidate guides and based on the secondary structure prediction choose the sgRNA-unc-22-1000 (were mfold (Fig.3.) predicts the correct pairings expected to occur between 5’variable and 3’constant regions, and therefore eliminates 5’ regions were ‘illegitimate’ intra-molecular hydrogen bonds might occur presumably preventing the proper interaction between tracrRNA and crRNA in the Cas9 complex).

Ours results provide the evidence for the sgRNA-unc-22-1000, guide RNA specifying the Cas9 cut prior to (NGG)n motif, were the second NGG is intended as PAM is fairly well tolerated when administrated as U6 driven transgene (with other components of transgenic CRISPR/Cas9). We foresee the other loci where repeat associated (NGG)n PAM could be identified would be valuable for experimental considering, given the multiple +3‘shifted’ guide RNAs could be predicted in one region. We speculate that the adjacent guide-internal and guide-external NGGs in the (NGG)n stretch could serve as auxiliary PAMs *in vivo* and perhaps could provoke the multiple Cas9 cuts.

Due to the particulars of the germline DNA transformation procedures in *C. elegans* (see discussion in (Arribere et al. 2015)), where DNA solution is injected into female part of the gonad of the parental animal, the Sp-Cas9 induced mutations in F1 expectedly occur in heterozygotic state and in case of recessive traits segregate in mendelian fashion phenotypically affected F2s (Friedland et al. 2013). In *C. elegans* however certain (<u>h</u>igh <u>i</u>ncidence of <u>m</u>ales) *him-6* alleles were demonstrated by (Haack & Hodgkin 1991) to segregate unusual progenies were both homologues of any given chromomosome derive from same gamete. We thought we therefore would test if, if this unusual property could be advantageous in CRISPR/Cas9 experiments in *C. elegans*. For those experiments we have chosen the CB1138 strain carrying weak recessive allele in *him-6* gene, reported as non-complementing with e1423 (Table 1, Table 16 Hodgkin et al. 1979). Weak *him-6* allele e1104 in CB1138 was expectedly preferred over e1423 (due to almost five-fold lower reported frequency of inviable zygotes (Table 2. and Table 16. in Hodgkin et al. 1979) presumably due to an excessive aneuploidy) for the DNA transformation procedures. The first experiment confirmed e1104 was successfully transformed with sgRNA-unc-22-1000 and Cas9 components, and that only F2 segregated desired unc22(−) phenotype, as it would be expected for the him-6(+). In the second experiment however (where sgRNA-unc-22-1000 was administrated as plasmid, expectedly increasing about ten fold the relative dose of U6 promoter driven guide RNA gene when compared to first experiment conducted with diluted PCR product), in the progeny of injected P0, we have observed and isolated the exceptional and fertile F1 unc-22(−) mutant. This observation, while certainly promising and desired could indeed reflect the scenario, where the exceptional euploid progeny with two maternally derived chromosomes IV (both him-6 and unc-22 are linked on IV) were mutated by sgRNA-unc-22-1000/Cas9 complex in the female germline of the injected hermaphrodite. Sequencing of the homozygous progeny of the above F1 unc-22(−) him-6(−) animal, revealed two lesions (105bp and 61bp deletions), suggesting the exceptional F1 unc-22(−) him-6(−) was a compound heterozygote. One model would assume here, either maternal line of descent (as a result of the non-disjunction at IV). Alternatively, however egg and sperm derived copies of chromosome IV was mutated. In any case the identified unc-22(−) F1 compound heterozygote (unc-22 (-105bp/-61bp)) would represent bi-allelic effects of Cas9. Indeed this could be an alternative (while not mutually exclusive) explanation for the observed Cas9 effects on unc-22 polyglycine encoding trinucleotide repeat associated PAM. While aware, that clearly more systematic, dedicated studies need to be designed to conclusively propose the model sufficiently explaining the particulars of inheritance of Cas9 triggered lesions in *C. elegans*, we propose: i. either, the direct Cas9 effects on paternal (sperm derived) chromosome needs to be excluded ii. or in the absence of the other corroborating evidence, it is not itself inconceivable that the generalized chromosomal non-disjunction might occur at some appreciable frequency in *C. elegans* germline chromosomes exposed to transgenic gRNA/Cas9 genes iii. Possibly alternative models i.e. assuming bi-allelic effects resulting from abortive DNA repair needs to be excluded.

Considering ours positive experience with CB1138 *him-6* in practical terms, we further suggest that *him-6* functions impaired in e1104 allele appear dispensable for CRISPR-Cas9. This appear relevant and possibly testable while working with X linked loci (due to the occurrence of the hemizygous X0 males). On the other hand however the presumed occurrence of triplo-X hermaphrodites (as a background result of the events of generalized non-disjunction involving sex chromosomes) and perhaps some other, more subtle events - results of generalized non-disjunction, might complicate the interpretation of some CRISPR-Cas9 induced phenotypes i.e. dpy phenotypes.

We have subsequently implemented the unc-22(−) hypersensitive line CL4176 [carrying *dvIs27*integrated transgene (Link et al. 2003) under the control of the smg-1(cc546ts)] the property identified previously (Kapulkin et al. 2005, Kapulkin WJ unpublished observation) as a convenient readout background in further experiments with sgRNA-unc-22-1000/Cas9. CL4176 is regarded as the model for beta-amyloid aggregation established by the Link lab (Fonte et al. 2002). The *dvIs27* transgene is integrated on the X chromosome in CL4176. *dvIs27* transgene is labeled with rol-6(d) marker and therefore could be easily counter-selected by crossings with males.

CRISPR/Cas9 approach would be very useful when working with this particular model of the APP/beta-amyloid dependent forms of Alzheimer disease i.e. researching the otherwise known mutations who suppress the temperature-sensitive lethal phenotype of the beta-amyloid (Lopez 2009). This is due the presence of unlinked temperature-sensitive mutation in the smg-1 (cc546ts) I. smg-1 (cc546ts) allele was developed by the Fire lab as system for conditional transgene expression and is required to regulate the expression of the beta-amyloid transgene in CL4176. Homozygous smg-1(ts) animals are superficially WT, which creates particular obstacle when attempting to introduce candidate suppressor mutations into doubly homozygous *dvIs27; smg-1*(cc546ts) CL4176. We hypothesized that CRISPR/Cas9 could be efficiently and elegantly implemented in the above model to create homozygous mutations in the complex transgenic background of CL4176. We first confirmed that sgRNA-unc-22-1000/Cas9 induced unc-22(−) phenotypes could be recovered with improved frequency amongst clonal F2 populations of CL4176. Homozygous *dvIs27; smg-1(ts)* animals are of Roller phenotype rol-6(su1006) and remain mobile when grown at 16°C. Introduction of the homozygous mutation into unc-22 via CRISPR/Cas9 in the above background efficiently converts mobile rol-6(su1006) animals into severe and spastic uncoordinated (unc) phenotype who remains distinct as almost entirely immobile (Supplement Movie 2). We have established a proof-of-the-principle evidence that CRISPR/Cas9 system could be used to select for desired mutations i.e. suppressor alleles, in the complex modified background based on the sgRNA-unc-22-1000 phenotypes. While we are uncertain on the exact nature of the hypersensitive effects of Cas9 induced disruptions into unc-22 locus in the CL4176, we note that most prevalent deletion (−61bp), is a frame altering lesion expected to result in mis-translation from the incorrect frame until the nearest stop codon, which is found in the intron. We note that given the specific properties of the *smg-1*(cc546ts) in with (−61bp) deletion in the background, certain CRISPR-Cas9 truncated alleles might translate more efficiently in *smg-1* (cc546ts) than in wild-type and perhaps contribute to the overall phenotype. Unexpectedly, in the CRISPR-Cas9 experiments with CL4176 we have recovered two (homozygous and presumably independent events, data not shown) integrated lines, where expectedly transient extrachromosomal array presumably containing plasmids encoding for genes providing sgRNA-unc-22-1000 and Cas9, were apparently chromosomally integrated. We emphasize those background integration events were observed solely based on segregation of the transgenic transformation markers co-administrated as separate marker plasmids and slow growth at non-permissive temperature. We note specifically those unexpected background integration events could easily go unnoticed, (if the transformation markers would be excluded from the above experiments), and therefore pose another unforeseen factor complicating the interpretation of the *in vivo* CRISPR/Cas9 experiments in *C. elegans*.

While comparing the above two sets of experiments with sgRNA-unc-22-1000/Cas9, the another unexpected observation, was that the same (−61bp) deletion have been observed independently in progeny born from the injected parental animals of the genotype him-6(−) and the *dvIs27;* smg-1(ts) (which is him-6(+)). Provided the parental animals represent two easily distinguished and genetically distinct *C. elegans* lines, we concluded that some of the most prevalent NHEJ-like lesions reoccurred in ours experiments carried on different background genotypes injected with the same mixture of the sgRNA-unc-22-1000/Cas9 delivering plasmids. We think the observed mutational bias observed in gRNA/Cas9 experiments is an intriguing phenomena and it would be interesting if observed in case of guide RNAs other than sgRNA-unc-22-1000 i.e. sgRNA-unc-22-999 and sgRNA-unc-22-998 or sgRNA-unc-22-1001 and 1002 etc, shifting the expected +3, PAM onto adjacent NGG present in impure polyglycine encoding repeat in the unc-22 gene. From the above experiments alone (carried by single person over ca. 16 month), we could not conclude on the reasons for the observed bias. Clearly in *C. elegans*, *in vivo* the Cas9 double stranded DNA is cut mostly at +3, PAM position (assumed based on Jinek et al. 2012) is followed by additional events (resulting in indels and larger deletions presumably as NHEJ-like products and MMEJ-like products). Those could be the biased DNA repair response events constrained by the local, germline chromatin structure or perhaps some other events affected by other factors i.e. Cas9 endonucleolytic activity for example was proposed to act together with exonucleolytic activity which might have been more robust *in vivo*.

Considering further the recurrent pattern apparent in the overall allelic spectrum of mutational events we observed while testing the Cas9 effects specified by sgRNA-unc-22-1000 at the (NGG)n trinucleotide repeat region embedding PAM, in addition to NHEJ-like events we observed the significant fraction of small frame restoring lesions. The small frame restoring mutations occurred as (−9bp), (−12bp) and two types of (−15bp) deletions (Table 2.), in the *unc-22* region specified by sgRNA-unc-22-1000. We have interpreted mutations in this class as the result of MMEJ-like repair events, restoring the +2(NGG)n frame encoding for poly-glycine tandem repeat. This class of alleles, presumably resulting from MMEJ-like repair, contributed to ~50% of the mutations we described in the overall observed allelic spectrum. MMEJ is regarded as mutagenic DSB repair mechanistically distinct from two other repair pathways: homologous recombination (HR) and Ku-dependent non-homologous end joining (NHEJ) (reviewed in McVey & Lee 2008). Microhomology-mediated end joining (MMEJ) is regarded as an alternative repair pathway, and is thought to rely on the internal alignment of short homologous sequences flanking the DSB (Sfeir &Symington 2015). While designing the guide RNA-unc-22-1000, we hypothesized that glycine encoding +2(NGG)n trinucleotide repeat embedding PAM, could provoke the MMEJ-like repair operating in *C. elegans*, and that is exactly what we have observed. MMEJ-like repair resulting from double stranded Cas9 cut would than relay on the defective alignment of the (NGG)n/(CCN)n repeats resulting in the small frame restoring deletions. We have observed this as frequent class of alleles represented by set of the consecutive deletions, in size from (−9bp), (−12bp) and (−15bp) recovered in the progeny born from independently injected parental hermaphrodites. Based on this observation we suggest that the MMEJ-like process is involved in the repair of (NGG)n PAM embedding regions in *C. elegans*. MMEJ therefore needs to be taken into consideration while designing CRISPR/Cas9 experiments in the nematode. MMEJ-like repair have been implicated in solving the Cas9 induced DNA breaks in genetic model organisms other than nematodes i.e. species of laboratory telosts and amphibians (Hisano et al. 2015)(Nakade et al. 2014) and mammalian cultured cells (Sakuma et al. 2015) some computer algorithms predicting the possible deletion pattern have been developed (http://www.rgenome.net/mich-calculator/ (Bae et al. 2014)). We suggest programs such us developed by NBRP Medaka vwould be useful to validate the targets for microhomologies (<http://viewer.shigen.info/cgi-bin/crispr/crispr.cgi >). Depending on the ultimate objectives of the genome editor, in some circumstances designer choice might intend to avoid regions prone to +2(NGG)n frame restoring MMEJ-like repair (i.e. in experiments requiring gene ablation resulting in null alleles). In other experimental schemes however, where more subtle than null alleles alterations to gene-protein sequence are desired, alleles resulting from frame restoring MMEJ-like repair products could be proven informative.

Considering ours positive results, confirming the involvement of frame restoring MMEJ-like repair, in the broader perspective we propose the 'modules' encoding for poly-glycine in +2(NGG)n position, (that are found quite frequently in coding areas of the genomes) overlap with ‘permissive’ PAM sites required by Cas9 *in vivo*. We foresee those (NGG)n PAM embedding repeats could pose an interesting targets for the genome-to-protein engineering. Given our results indicate the +2(NGG)n poly-glycine encoding stretch could be cleaved by Cas9 and repaired by frame restoring MMEJ resulting in the removal of the glycine-repeat strech from the the mature protein. Glycine is a smallest encoded aminoacid incorporated into polypeptide chains and presumably the mature protein lacking the poly-glycine stretch could be regarded as 'constricted' (Fig.4.C). Clearly the poly-glycine encoding stretches could have different functions in context of different proteins, however ours approach permits for this type of engineering, bypassing the need for less efficient exogenous template directed genome-editing.

Finally, we have established that fraction of the Cas9 induced NHEJ events specified by sgRNA-unc-22-1000 co-occur with when sgRNA-DPY-10 and cn64 (Aribere et al. 2015) converting oligo are co-injected into hermaphrodite gonad. Provided we have isolated two unc-22(−) segregating lines out of seven rol or rol/dpy lines, we think this observation would deserve a comment. While we agree with others and think the NHEJ-like events are more frequent than repair events involving the co-administrated ssDNA oligonucleotide template, we note that dpy-10/cn64 co-conversion is more frequent than NHEJ-like events observed in (NGG)n type PAM repeat region of unc-22. This however might be explained by realizing that different guides might have very different optima *in vivo* and different efficiencies particularly if used at different concentrations. This might be due to the guide-intrinsic factors as well as guide-extrinsic factors i.e. guide-RNA toxicity and context sequence/ chromatin state dependent, respectively. The above concerns however do not change ours overall conclusions that co-conversion could be observed as co-occuring in sgRNA-unc-22-1000 guide-RNA mixing experiments, efficiently demonstrating robust co-conversion from the synthetic oligonucleotide template at the unlinked dpy-10 locus. We further suggest that dual selection might be advantageous in enriching for the rare or otherwise inefficient Cas9 events.

## Acknowledgments

We thank the MPI-CBG Dresden for the hostile environment to conduce the experiments. In particular we grateful for professor Anthony Hyman to host us, and providing the postdoctoral stipend to WJK. We are grateful to colleagues in the Hyman lab in particular to Susane Ernst for providing the supporting atmosphere and other colleagues in the Hyman lab. Other collegues at MPI, in particular Mark Leaver (Hyman lab) and Christian Eckmann (MPI-CBG) for correcting and carefully suggesting on the initial draft of the manuscript submitted to WBG. We acknowledge support from members of the MPI-CBG facility services: Stephanie Marret, Dana Suchold, Stephan Janosch, Sussane Hasse, and Andrey Pozniakowski (Hyman lab). DNA sequencing and informatic facilities of MPI-CBG are acknowledged accordingly for providing the excellent environment expertise and providing the accurate measurements and DNA analyses of the CRISPR-Cas9 alleles. We are indebted to colleagues providing us with recombinant DNA constructs via Addgene. We are eternally grateful to Andrew Fire (Stanford) for sharing the enthusiasm over the CRISPR system, insightful and inspiring discussions and constructive comments and suggestions on the manuscript.

## Notes added in proof

Initial description of ours experiments have appeared in Worm Breeders Gazette (2014) <wbg.wormbook.org/2014/10/20>.

During the time ours manuscript was prepared for publication, (Farboud and Meyer 2015) working with several different guides targeting sequences loci other than UNC-22 reported dramatic enhancement of genome editing with Sp-Cas9 with improved consensus. We note ours results with repeat-associated PAM within unc-22 polyglycine encoding repeat are consistent with their findings. Also, within the time ours manuscript was prepared for submission the other group reported experiments with different guide RNA within unc-22 gene. We note that since the CRISPR-Sp-Cas9 improvement (Jinek et al. 2012) were implemented into *C. elegans* field (Friedland et al. 2013) only a handful set (i.e. Kim H et al. 2014) of all possible of more than thousand different candidate guides predicted to lay within unc-22 exonic segments, were tested *in vivo*, until this manuscript was finally prepared (Warsaw, May 14 2016).

